# Dynamic prioritisation of sensory and motor contents in working memory

**DOI:** 10.1101/2024.11.12.623203

**Authors:** Irene Echeverria-Altuna, Sage E.P Boettcher, Freek van Ede, Anna C. Nobre

## Abstract

Internal selective attention prioritises both sensory and motor contents in working memory to guide prospective behaviour. Prior research has shown how attention modulation of sensory contents is flexible and temporally tuned depending on access requirements, but whether the prioritisation of motor contents follows similar flexible dynamics remains elusive. Also uncharted is the degree of co-dependence of sensory and motor modulation, which gets at the nature of both working-memory representations and internal attention functions. To address these questions, we independently tracked the prioritisation of sensory and motor working-memory contents as a function of dynamically evolving temporal expectations. The design orthogonally manipulated when an item location (left vs right side) and associated prospective action (left vs right hand) would be relevant. Contralateral modulation of posterior alpha (8-12 Hz) activity in electroencephalography (EEG) tracked prioritisation of the item location, while contralateral modulation of central mu/beta (8-30 Hz) activity tracked response prioritisation. Proactive and dynamic alpha and mu/beta modulation confirmed the flexible and temporally structured prioritisation of sensory and motor contents alike. Intriguingly, the prioritisation of sensory and motor contents was temporally uncoupled, showing dissociable patterns of modulation. The findings reveal multiple modulatory functions of internal attention operating in tandem to prepare relevant aspects of internal representations for adaptive behaviour.

## Introduction

Internal selective attention selects and prioritises working-memory contents according to goals, expectations, and other control-related cognitive processes (Myers et al., 2017; Nobre, 2018; Nobre & van Ede, 2023; Oberauer, 2019; Oberauer & Hein, 2012; van Ede & Nobre, 2023). Far from changing randomly, our environment is structured across space, time, and other attributes (e.g., Nobre & van Ede, 2018; Schwartz et al., 2007). Internal attention can use the expectations emerging from these regularities to prioritise working-memory contents and, thereby, proactively guide ongoing behaviour.

Numerous studies have found that expectations about the sensory qualities of objects (i.e., location, colour, and other stimulus features) can proactively guide internal attention to the relevant working-memory contents thus facilitating subsequent task performance (Griffin & Nobre, 2003; Landman et al., 2003; Lewis-Peacock et al., 2012; Oberauer & Hein, 2012; Serences et al., 2009; for reviews see Souza & Oberauer, 2016; van Ede & Nobre, 2023). A growing literature has also highlighted the pragmatic nature of internal attention, demonstrating that it can proactively prioritise action-related contents in working memory (Boettcher et al., 2021; Formica et al., 2021; González-García et al., 2020; Henderson et al., 2022; Kikumoto et al., 2022; Nasrawi et al., 2023; Nasrawi & Van Ede, 2022; Rösner et al., 2022; Schneider et al., 2017; van Ede et al., 2019a). Sensory and action-related contents can co-exist in working memory. When a working-memory item is probed, selection of its sensory- and motor-associated contents starts concurrently (van Ede et al., 2019a). Beyond this initial demonstration, the dynamics of the prioritisation of sensory and motor contents that co-exist in working memory remain elusive.

Electroencephalography (EEG) provides a powerful method for independently tracking the prioritisation of sensory- and action-related contents in working memory. The selection of action plans in working memory is mirrored by a relative reduction in mu/beta (8-30 Hz) activity contralateral to the prospective action hand at central electrodes (e.g. Kaiser et al., 2001; Pfurtscheller et al., 2000; Schneider et al., 2017; van Ede et al., 2019a). In turn, the selection of visual representations held within the spatial layout of working memory is mirrored by a relative reduction in contralateral alpha-frequency (8-12 Hz) activity in posterior electrodes (e.g., Mok et al., 2016; Myers et al., 2014; Poch et al., 2014; Schneider et al., 2016; van Ede et al., 2019a; van Ede et al., 2017; Wallis et al., 2015).

It is increasingly clear that the prioritisation of sensory contents in working memory is highly flexible and dynamic. For example, selecting a sensory item in working memory does not necessarily compromise the representation of other competing items, illustrating the flexible and reversible nature of working-memory content prioritisation (Christophel et al., 2018; De Vries et al., 2017, 2018; LaRocque et al., 2013; Lepsien & Nobre, 2007; Lewis-Peacock et al., 2012; Muhle-Karbe et al., 2021; Myers et al., 2018a; Rerko & Oberauer, 2013; van Ede et al., 2021; Van Moorselaar et al., 2015). Moreover, the prioritisation of sensory contents, as measured with EEG activity modulations or pupil size, changes according to temporal expectations concerning the likely time of an item to be probed (De Vries et al., 2018; Jin et al., 2020; van Ede et al., 2017; van Loon et al., 2017; Zokaei et al., 2019). In contrast, it remains unknown whether the prioritisation of action-related working-memory contents can be similarly reversible and tuned to the temporal structure of the task. Additionally, while sensory- and action-related working-memory contents can be prioritised concurrently (van Ede et al., 2019a), it is unclear whether these two aspects of the representation are functionally bound. Are they two sides of the same integrated internal representation and, therefore, does their prioritisation co-evolve in lockstep according to the dynamic structure of the task? Or are sensory and motor contents in working memory dissociable and capable of separate modulation?

To address these questions, we designed a task that zoomed into how temporal expectations guide the dynamic prioritisation of sensory- and action-related contents in working memory. Item location (left vs right side) was orthogonally crossed with required action (left vs right hand). The design thereby enabled tracking the prioritisation of sensory contents (location) and prospective actions independently (see also Boettcher et al., 2021; Nasrawi et al., 2023; van Ede et al., 2019a). Contralateral-vs-ipsilateral posterior alpha (8-12 Hz) and central mu/beta (8-30 Hz) modulations provided markers of the prioritisation of sensory- and action-related contents in working memory, respectively. To foreshadow the results, the prioritisation of both sensory- and action-related working-memory contents evolved flexibly as a function of dynamically changing expectations. However, their prioritisation was not continuously temporally coupled, suggesting that multiple modulatory functions can operate in tandem – and develop independently – on different aspects of working-memory representations.

## Methods

### Participants

This study was approved by the Central University Research Ethics Committee of the University of Oxford (R57489/RE006). The sample size of the present study was based on previous studies investigating related questions (Boettcher et al., 2021; van Ede et al., 2019a). Thirty-one volunteers participated in two experimental sessions, which took place on different days. All participants self-reported having normal or corrected eyesight. They provided written consent before each session and were reimbursed at £15/h for their participation.

Exclusion criteria were pre-defined. Participants were excluded from further analyses if their average performance error or reaction time (RT) was above 3 standard deviations (SDs) from the mean error or RT, respectively, across all participants in either session. Data from one participant were excluded from further analyses based on these criteria. The final sample (n = 30) had an average age of 23.57 (SD: 3.85). Three individuals were left-handed and twenty-seven were right-handed by self-report; six participants identified as male, twenty-three as female, and one as non-binary.

### Experimental procedure and stimuli

We designed a visual-motor working-memory task to study the flexible and dynamic prioritisation of visual and action-related contents in working memory (**Figure 1a**). Participants were shown two coloured, tilted bars and were asked to report the tilt of one of the bars at the end of each trial. Retro-cues that matched or did not match the colour of one of the encoded bars were informative or noninformative regarding the item participants had to report at the end of the trial, respectively. Participants reported the orientation of one of the encoded bars at the end of each trial. Importantly, bar orientation (left vs right) was linked to the response hand (left vs right). Additionally, the location of the bars on the screen (left vs right) and the direction of their orientation (left vs right) were manipulated orthogonally. Therefore, the prioritisation of bar locations and prospective hand actions could be tracked independently using their respective EEG markers (see also Boettcher et al., 2021; Nasrawi et al., 2023; van Ede et al., 2019a; van Ede et al., 2019b).

**Figure 1.**
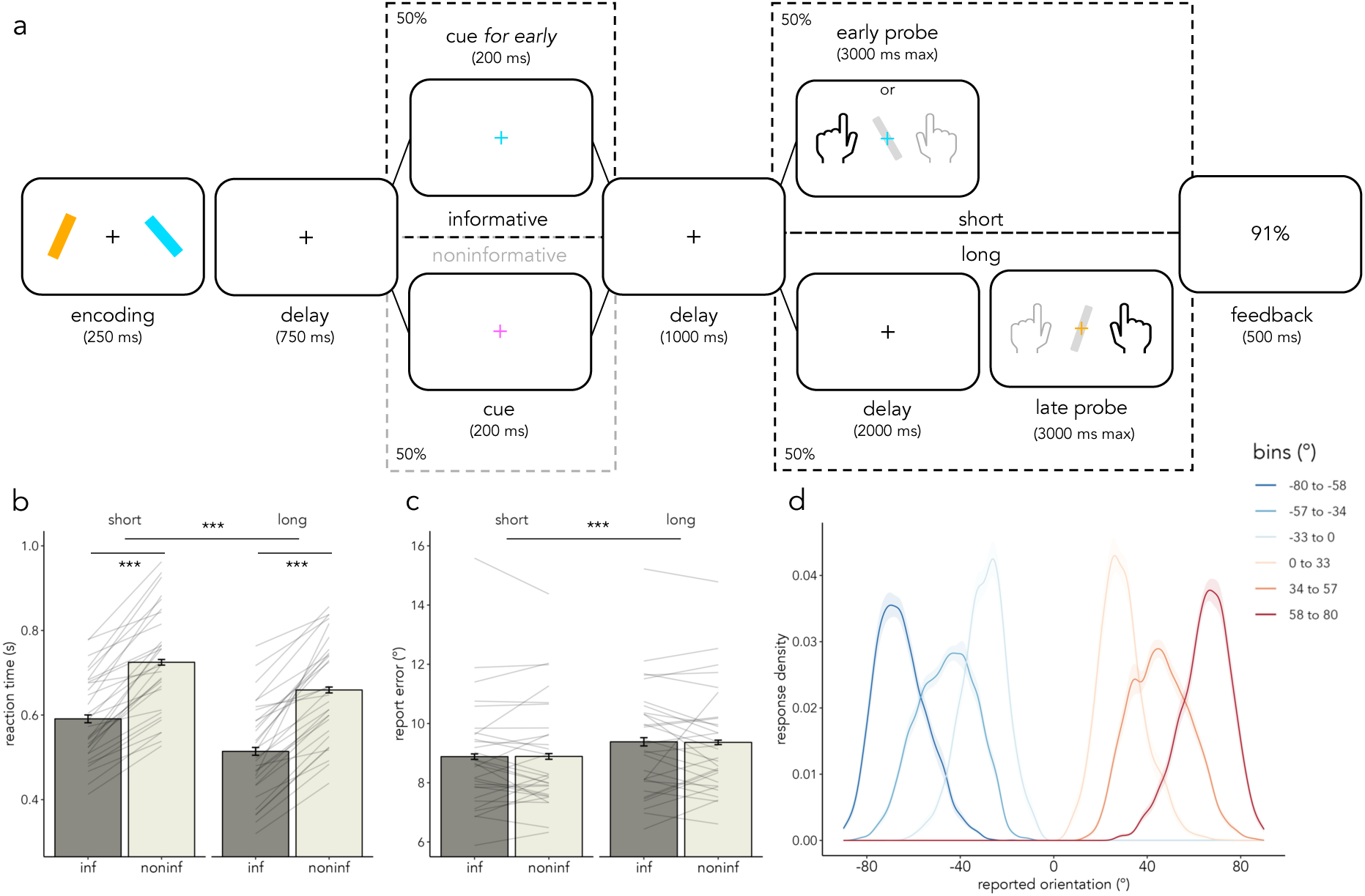
Task design and behavioural results. a) Trial schematic. Two coloured, tilted bars (one on the left and the other on the right, one tilted to the left and the other to the right, with location and tilt being orthogonally manipulated) were displayed at encoding. In half of the trials (informative) a retro-cue matching the colour of one of the two bars was shown; in noninformative trials (50%), a cue with a different colour appeared instead. In informative trials, if the delay after cue offset was short (1 s), participants were probed about the cued item. Alternatively, if the delay was long (3 s), they had to report the other (uncued) item. In noninformative trials, the retro-cue did not match the colour of either bar and the probed item was unpredictable. Reaction time (s) (panel b) and report error (°) (panel c) are plotted in informative and noninformative and short and long trials. Thin grey lines represent individual participants, error bars represent the SEM, light grey represents noninformative trials, and dark grey depicts informative trials. d) Average response density as a function of the reported orientation and the orientation of the reported bar. Shaded areas represent SEM.

The experimental script was generated using the Psychophysics Toolbox version 3.0.18 (Brainard, 1997) on Matlab 2022a (The Mathworks Inc., Natick, NA, USA). Participants sat in a dimly lit room, ∼60 cm away from a monitor (Dell U2312HM; 1920×1080 pixels resolution; 100-Hz refresh rate).

The display background was grey (#7F7F7F) for the duration of the task. At the beginning of each trial, participants saw the encoding display made of a white central fixation cross (#FFFFFF; 14 pixels) and two coloured, tilted bars shown for 250 ms. One bar was always on the left and the other on the right side of the screen. Each bar was centred 192 pixels (5.2 DVA) away from the central fixation cross, had a length of 192 pixels and a width of 38 pixels. One bar was always tilted leftward and the other rightward. Bar tilts were drawn from one of three possible bins with a uniform distribution of orientations (+/-10-33°, +/-34-57°, and +/-58-80°). To avoid cardinal effects, none of the displayed bars had orientations along the vertical or horizontal meridians. Orientation bins were counterbalanced so that all bins were equiprobably sampled for each participant. Bar location and tilt were orthogonally manipulated, such that leftward- and rightward-tilted bars were equally likely to appear on the left or right side of the screen across trials. The two bars on the encoding display had two different colours out of four possible, highly distinguishable colours: light blue (#00DEFF), orange (#FFAC00), pink (#FF62FF), and green (#00ED82). Colours were counterbalanced across trials such that all possible combinations of colour, location, and tilt were displayed.

Following the encoding display, the coloured bars disappeared, and the central fixation cross remained on the screen for 750 ms (first delay; **Figure 1a**). Subsequently, the colour of the central fixation cross changed for 200 ms (retro-cue). In half of the trials (*informative;* 50%), the colour of the retro-cue matched the colour of one of the two bars displayed previously. In the other half of the trials (*noninformative;* 50%), the fixation cross changed to a different colour that matched neither encoded item on that trial. The cue colour was chosen from the four possible colours detailed above. Cue colour was counterbalanced such that all colours acted as cues in the same number of trials across each block of the task and appeared equiprobably in informative and noninformative trials.

In half of the informative and half of the noninformative trials, the response probe appeared 1 s after cue offset (*short* trials; 50%). In the other half of the trials, the response probe appeared 3 s after cue offset (*long* trials; 50%). During the post-cue delay (short and long), only a white, central fixation cross remained on the screen.

Crucially, in informative trials, the combination of two pieces of information predicted which of the two bars would be probed at the end of the trial with 100% validity: 1) the colour of the retro-cue and 2) the duration of the delay between the cue and the probe. When the cue was informative (matching the colour of one of the two bars displayed at encoding) and the delay following the cue was short (1 s), participants were prompted to report the orientation of the cued bar at the end of the trial (100% validity). Alternatively, in informative trials with a long delay (3 s), participants were always required to report the orientation of the other (uncued) bar (100% validity). Therefore, participants were encouraged to attend to the retro-cue and to track the duration of the subsequent delay to know which of the two bars they would be asked to report. In informative trials, they could anticipate reporting the item with the cued colour after the short interval. Once the short interval passed, they could shift the focus of attention to the item with the other colour. In noninformative trials, which of the two bars would be probed was unpredictable to participants.

Following the delay, the colour of the central fixation cross changed again (probe) prompting participants to reproduce the tilt of the colour-matching bar (**Figure 1a**). To this end, participants used the “F” and “J” keys on the keyboard with their left and right index fingers respectively. Upon response initiation, a grey bar appeared centrally in the vertical position. Pressing the “F” key led to a counterclockwise rotation of the grey bar and pressing the “J” key made it move clockwise. Participants were instructed to release the key when the grey bar reached the desired orientation. The size of the grey bar was the same as that of the coloured bars on the encoding display. The grey bar did not rotate further than 90° in either direction. Therefore, leftward tilted bars could only be reported with the left hand and rightward tilted bars with the right hand. The response key could not be changed once a response was initiated. Participants were instructed to report the orientation as quickly and accurately as possible. They had a maximum of 3 s to respond from probe onset.

If participants responded with the correct key (F for leftward-tilted bars and J for rightward-tilted bars), they received visual feedback about the percentage accuracy of their response. If they pressed the incorrect key, they received the message “Wrong target!”. If they did not respond, they saw “Too slow!”. All feedback appeared centrally in white letters and was displayed for 500 ms. Inter-trial intervals (ITIs) followed a beta distribution with a minimum of 3 s, maximum of 10 s, and average of 5.5 s. During each ITI, a white fixation cross was displayed centrally.

The present study was divided into two separate sessions in which participants repeated the same task procedure on different days. The two visits allowed participants to become familiar with the task instructions and procedures. The first visit consisted of a behavioural session in which participants completed eight blocks of 32 trials each for ∼1 h. The second visit consisted of an EEG and eye-tracking session in which participants completed 15 blocks with 32 trials each for ∼2.5 h. In the EEG session, participants could rest for 10-15 minutes after every 5 blocks. Before beginning the first session of the experiment, participants were walked through the task instructions and asked to practice the procedure for at least one block.

### Behavioural data analysis

Behavioural data were analysed using the R statistical programming language (version 4.2.1; R Core Team, 2021) and R studio (version RStudio 2022.07.1; RStudio Team, 2022). Report errors were calculated as the absolute difference between the reported orientation and the orientation of the probed bar. RT was defined as the time from probe onset until response initiation. Trials without a response, trials with RTs faster than 100 ms, trials with RTs slower than three times the SD of the average RT per participant, and trials with report errors higher than three times the SD of the participant-averaged error were excluded from further behavioural analyses. This resulted in an average removal of 2.38% (SD: 0.72%) of trials in the behavioural session and 2.82% (SD: 0.6%) in the EEG session. The average RT and report error per participant and per condition were calculated for each session. Given the equivalent behavioural pattern across sessions (**Supplementary Tables 1-4**), RT and report errors from both sessions were collapsed (**Figure 1b,c**). Additionally, we calculated the average response density as a function of the reported tilt and the tilt of the probed item (**Figure 1d**).

The statistical significance of the dependent variables of interest across participants (report error and RT) was tested using 2 x 2 repeated-measures analysis of variance (ANOVA) with informativeness (informative and noninformative) and duration (short and long) as factors. The metric of effect size was **η**^2^ and the within-subject standard error of the mean (SEM) was quantified using the normalised data (Morey, 2008).

### EEG: acquisition and preprocessing

EEG was acquired using a 64-channel Quik-Cap Neo Net cap (Ag/AgCl electrodes), Synamps amplifiers, and the CURRY 8 acquisition software (Compumedics Neuroscan). Sixty-four channels were distributed across the scalp following the international 10-10 positioning system. Data were referenced online to a reference channel positioned between Cz and CPz. Another channel (AFz) was used as the ground. Vertical and horizontal electrooculograms (EOG) were simultaneously recorded using a bipolar system integrated into the cap. Horizontal EOG electrodes were placed on the side of each eye and vertical EOG was positioned above and below the left eye. The electrocardiogram (ECG) was measured with an integrated, bipolar set-up. The upper ECG was placed on the left ribcage, centred above the left chest; the lower ECG was placed on the side of the left ribcage. During set-up, electrode impedance was lowered to below 5 kiloohms where possible and was at least below 10 kiloohms. Data were digitised at 1,000 Hz and filtered at 500 Hz during acquisition.

All EEG data were processed and analysed in Python 3.11.9 using MNE-Python (version 1.5.1; Gramfort, 2013) and custom-made scripts. First, data were notch-filtered at 50 Hz, and high-pass (0.05 Hz) and low-pass (40 Hz) filtered with a Finite Impulse Response (FIR) filter. Subsequently, the continuous data were visually inspected. Channels deemed “bad” were interpolated using spline interpolation as implemented with the *interpolate_bads* function in MNE-Python. Then, data were downsampled to 250 Hz and re-referenced to the sensor average. An independent component analysis (ICA) identified artifacts related to eye movements and heart-related activity by correlating individual ICs with the EOG and ECG signals. Based on the correlation values and visual inspection of the IC time courses and topographies, ICs capturing eye- and heart-related activity were identified and subtracted from the EEG data. An average of 2.87 ICs (SD: 0.8; range: 1-5) were removed per participant.

Subsequently, the raw EEG data were epoched around cue onset and probe appearance (−0.25 s to 1.25 s in short trials and −0.25 s to 3.25 s in long trials), and activity during the baseline period (−0.25 to 0 s) was subtracted from each epoch. A surface Laplacian transform was applied to the epoched data using the *compute_current_source_density* MNE-Python function to reduce the effects of volume conduction and thus increase the spatial resolution and interpretability of the results (see also van Ede, et al., 2019a).

Trials included in the EEG analysis were limited to those in which participants had pressed the correct key and RT was higher than 100 ms. Noisy trials, as identified on the epoched EEG time courses using a Generalized ESD procedure (Rosner, 1983), were also excluded. In total, 89.67% (SD: 5.34%) of trials were used in subsequent analyses.

### EEG: time-frequency analyses

All EEG analyses focused on the period of interest from retro-cue onset until probe onset (see **Figure 1a**). The retro-cue in informative trials was hypothesised to prompt participants to direct attention to the cued sensory (location) and action-related (response hand) contents in working memory. Importantly, it was hypothesised that in long trials, participants would “shift” to prioritising the other stimulus location and response hand after the short interval lapsed. Crucially, given the orthogonal manipulation of bar location and tilt, prioritisation of locations and action plans could be tracked independently with EEG (see also Boettcher et al., 2021; Nasrawi et al., 2023; van Ede et al., 2019a). Specifically, it was hypothesised that EEG alpha-frequency activity (8-12 Hz) at occipital electrodes contralateral to the location of the prioritised item would be modulated. In parallel, it was predicted that EEG mu/beta activity (8-30 Hz) at central electrodes contralateral to the prioritised response hand would be modulated. Alpha-band activity (8-12 Hz) at left (PO7) and right (PO8) visual sensors was used to track sensory-related prioritisation. Mu/beta activity (8-30 Hz) in left (C3) and right (C4) motor electrodes was used to track action-related prioritisation (see also Boettcher et al., 2021; van Ede et al., 2019a). For confirmation purposes, the same analyses were performed using a larger cluster of lateralised visual and motor sensors (**Supplementary Figure 1**).

The epoched EEG time series were transformed into their time-frequency decompositions using the SAILS Python toolbox (Quinn & Hymers, 2020; Quinn et al., 2021). This was done by convolving the time series with a complex 3-cycle Morlet wavelet from 2 to 40 Hz in steps of 1 Hz. The 50 ms around the two edges of each epoch were subsequently cropped to remove any edge artefacts related to the time-frequency decomposition. Time-frequency activity in lateralised visual (PO7/PO8) and motor (C3/C4) channels was contrasted between trials in which either the stimulus location or the prospective action hand (related to the tilt), respectively, was contralateral vs ipsilateral to each channel. This was calculated and normalised as follows: [((contra – ipsi) / (contra + ipsi)) * 100], separately for left and right sensors per participant. Subsequently, the contrast across both sides was collapsed in the visual and motor selection conditions separately and averaged across participants.

The same procedure was followed with each symmetrical electrode pair to create the topographical lateralisation maps shown in **Figure 2e,f**. The topographies depict the right-vs-left contrasts at the time points corresponding to the statistically significant clusters in the panels above (**Figure 2a,b**; see **Cluster-based permutation testing** for details).

**Figure 2.**
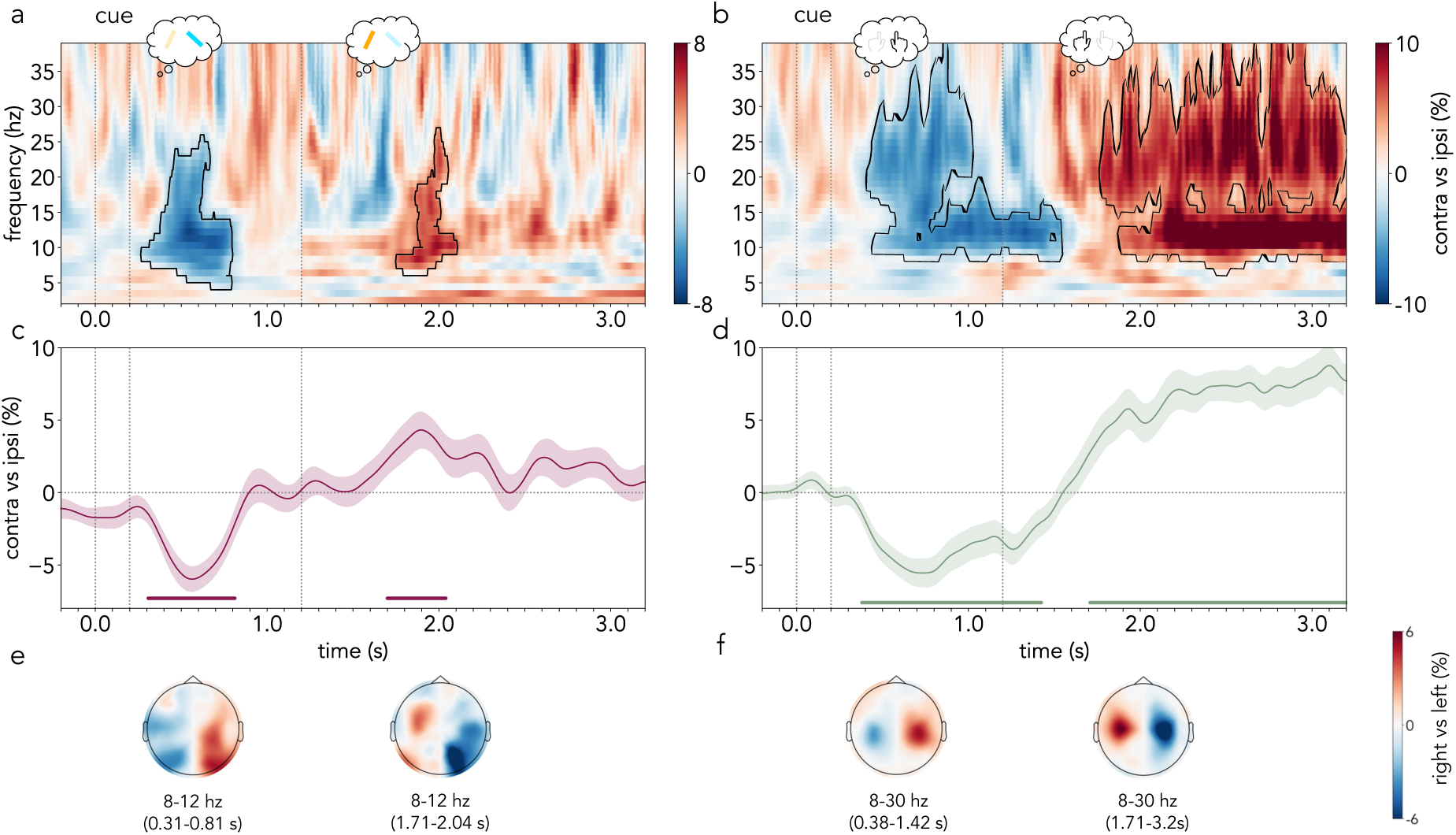
Lateralised frequency-specific EEG activity locked to cue onset. a) Contrast between EEG time-frequency activity contralateral vs ipsilateral to the cued bar location in occipital sensors (PO7/PO8) divided by summed contralateral and ipsilateral activity (expressed as a percentage) in informative trials vs noninformative trials. b) Contrast between EEG time-frequency activity contralateral vs ipsilateral to the cued prospective action in central sensors (C3/C4) divided by summed contralateral and ipsilateral activity (expressed as a percentage) in informative trials vs noninformative trials. c,d) Cross-participant average alpha (8-12 Hz; c) and mu/beta (8-30 Hz; d) activity difference between contralateral and ipsilateral sensors to the cued location and action, respectively, in informative trials. e,f) Topographies represent the average frequency-specific activity in right-vs-left contrasts in informative trials across all sensor pairs during the time-windows which correspond to the alpha clusters in panel c or mu/beta clusters in panel d, respectively. Black outline in time-frequency spectra indicates statistically significant clusters. Shaded areas represent the SEM and cluster-permutation corrected significant time points are indicated with horizontal lines. The first part of the time-frequency spectra in panels a and b and of the time course in c and d (−0.2–1.2 s) corresponds to the average of short and long trials, and the second part (1.2–3.2 s) corresponds to long trials only. The vertical dotted lines represent (from left to right) the onset (0 s) and offset (0.2 s) of the retro-cue and the time of probe appearance in early trials (1.2 s).

Additionally, to generate the time course of posterior alpha and central mu/beta modulations, signals from the pre-defined frequencies (alpha: 8-12 Hz; mu/beta: 8-30 Hz) were averaged at the selected channels **(Figures 2b,d)**. To increase sensitivity and improve visualization, the trial-averaged alpha and mu/beta time courses for each participant were smoothed using a Gaussian kernel with a standard deviation of 40 ms (see also van Ede et al., 2019a).

To increase their comparability, the alpha and mu/beta modulation time courses were each scaled to range between −1 and 1 per participant. Subsequently, we used cluster-based permutation (see below) to statistically compare the alpha and mu/beta time courses across participants (**Figure 3a**). The average time-frequency activity in the early period (−0.2 to 1.2) includes both short and long trials. The later period (1.2 to 3.2) includes only long trials.

**Figure 3.**
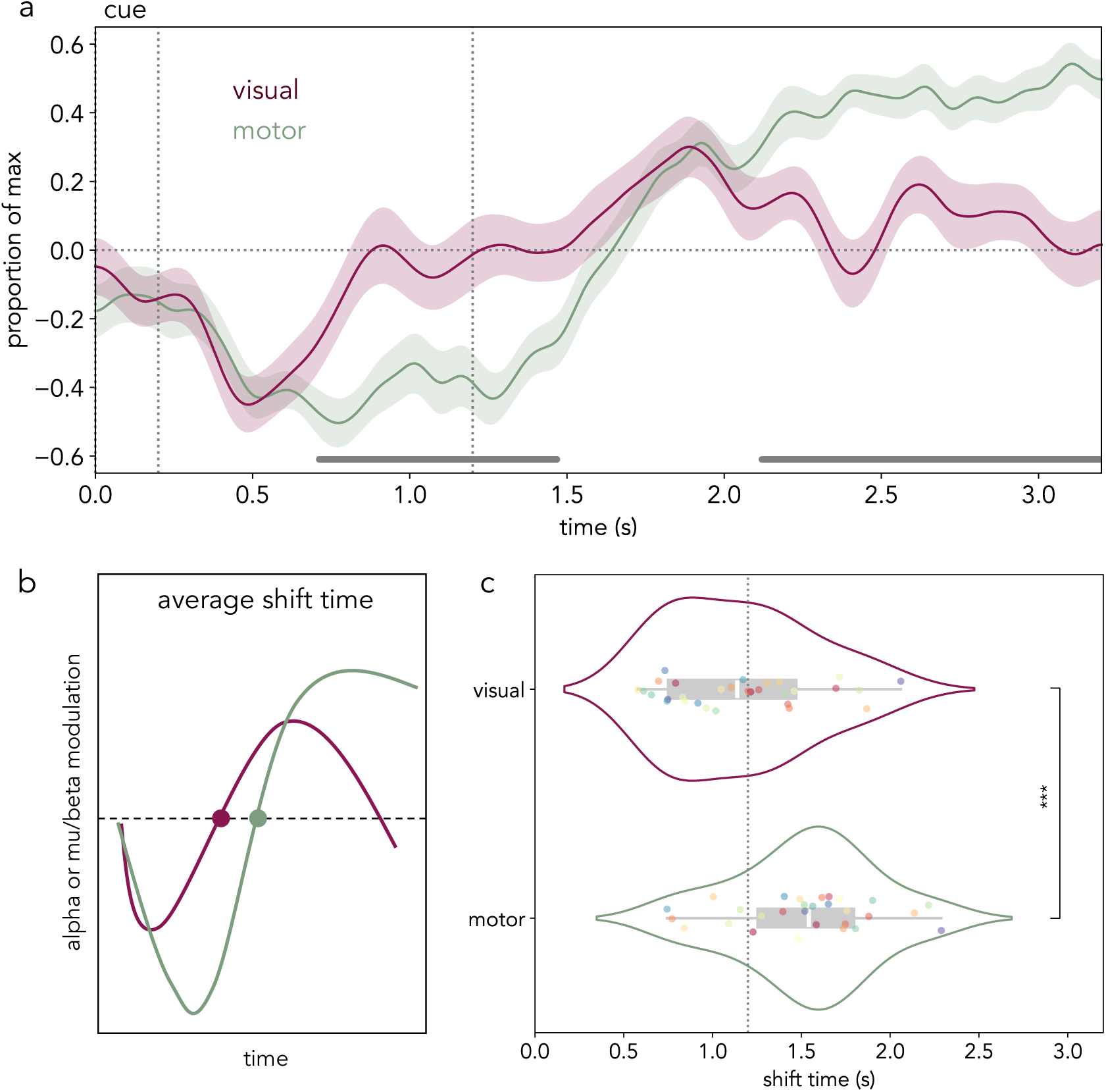
The alpha and mu/beta modulation time courses do not co-evolve in lockstep. a) Average alpha (8-12 Hz) activity difference between contralateral and ipsilateral sensors to the cued location (burgundy) and average mu/beta (8-30 Hz) activity difference between contralateral and ipsilateral sensors to the cued action (green) scaled to range between −1 (minimum) and 1 (maximum) per participant. Horizontal grey lines depict time points that were significantly different between the normalised alpha and mu/beta modulation time courses in a participant-wise cluster-based permutation test. Shaded areas represent the SEM. The vertical dotted lines represent (from left to right) the offset (0.2 s) of the retro-cue and the time of probe appearance in early trials (1.2 s). b) Schematic representation of the visual and motor shift time calculation procedure. For each participant, we calculated the minimum and maximum points of alpha and mu/beta modulation and estimated the average zero-crossing time (shift time; see *Methods*). c) Violin plots depicting the distribution of visual (top) and motor (bottom) average shift times across participants. Coloured dots represent individual participant mean values, the white line inside the grey box represents the mean, and the edges of the grey box represent the first (left) and third (right) quartiles. Statistical significance is depicted with asterisks.

### EEG: visual and motor shift time quantification

Next, the temporal relation between the time courses of the two frequency bands of interest (alpha and mu/beta) was investigated. With this aim, a set of key control points was identified on the trial-average time courses. First, the participant-averaged alpha and mu/beta time courses were smoothed using a Gaussian kernel with a standard deviation of 120 ms to facilitate the identification of the control points. To enhance comparability between the signals, the alpha and mu/beta time courses were scaled to range between −1 and 1, respectively. For each participant, the minimum point of alpha and mu/beta attenuation was identified within a pre-defined time window (0.1 to 1 s from cue onset), and the maximum point of alpha and mu/beta was identified in a later window (1 to 2.5 s from cue onset). Subsequently, the average time of zero crossing (i.e., lateralisation reversal) between the identified minimum and maximum points was identified as the average shift time, as depicted in **Figure 3b**. When participants had multiple zero crossings, the average shift time was calculated. When participants had a single zero crossing, we considered the latter to be the average shift time. A two-sided paired-samples t-test compared the average shift times of the alpha and the mu/beta time courses across participants (**Figure 3c**). The effect size was reported as Cohen’s d.

Next, we followed a bootstrapping procedure in which half of the trials in each participant were randomly sub-sampled over 100 iterations (**Figure 4a**). For every subset of trials from a single participant, the contralateral-versus-ipsilateral alpha and mu/beta time courses were calculated, smoothed, and scaled as detailed above. Subsequently, the control points were identified. Next, we tested for any existing correlations between the mean visual shift time and the mean motor shift time across all iterations using a linear mixed-effects model (LMM; using the *lmer* from the R package *lme4*) that incorporated participant-related random effects as follows:

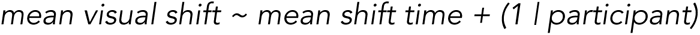

**Figure 4.**
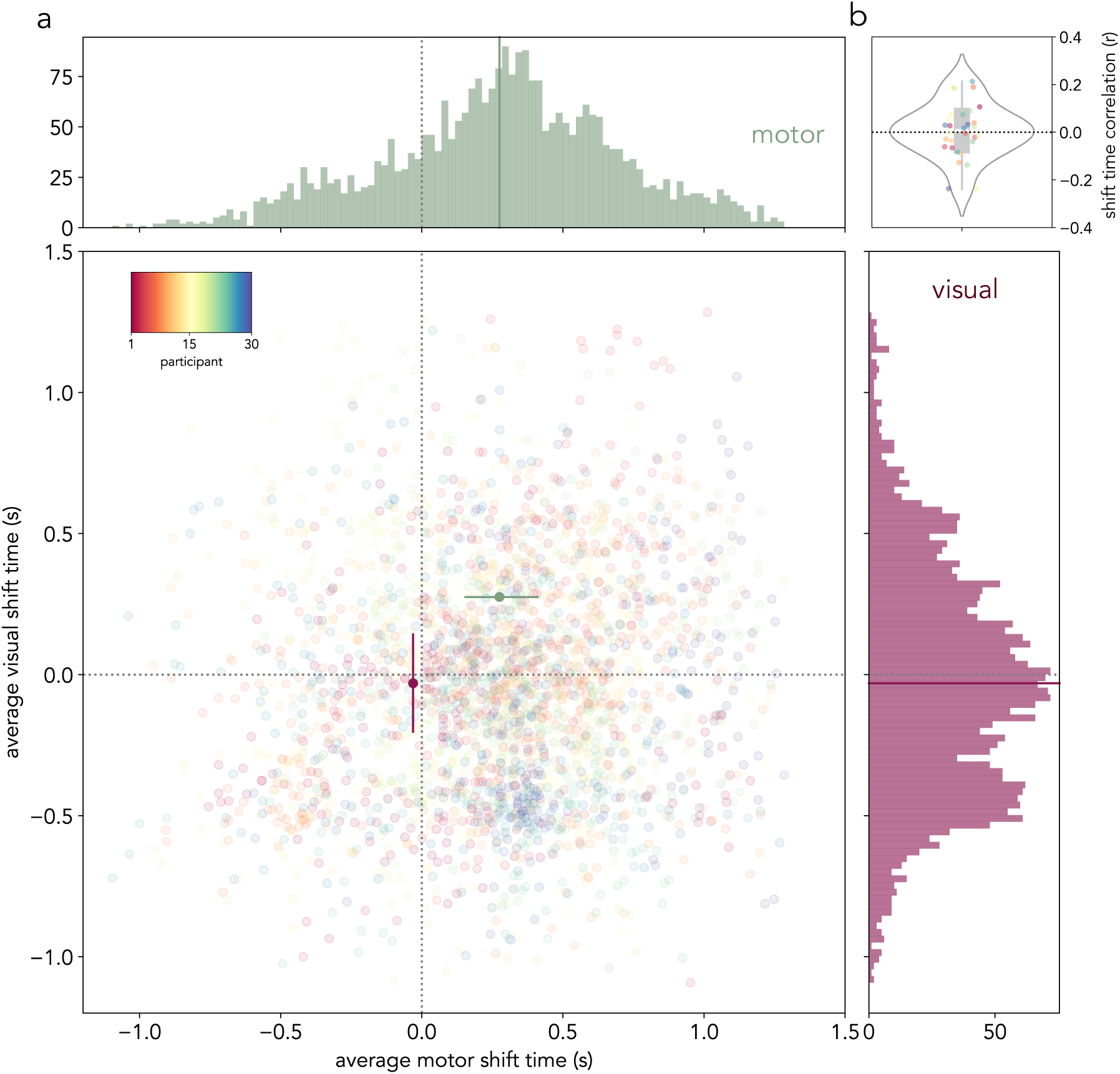
Lack of correlation between the shift times of the alpha and mu/beta modulation time courses. a) Top and right: the histograms represent the distribution of average mu/beta (green) and alpha (burgundy) shift times, respectively, on all the iterations of the bootstrapping procedure (50 bins per histogram). Centre: scatter plot of the average mu/beta and alpha shift times in each of the 100 bootstrapping iterations across all 30 participants as plotted on the mu/beta (x-axis) and alpha (y-axis) time space. Both axes are centred around the time of early probe onset. Vertical and horizontal dotted lines represent the time of early probe appearance (1.2 s after retro-cue onset). The burgundy circle represents the participant-averaged shift time as estimated on the alpha time course, and the green circle represents the shift time estimated on the mu/beta activity average. The lines on left and bottom the green and burgundy circles, respectively, depict the participant-averaged minimum motor and visual shift times, respectively. The lines on right and top the green and burgundy circles, respectively, depict the participant-averaged maximum motor and visual shift times, respectively. The colours of the scattered dots depict individual participants. b) Pearson’s r values of the correlation between motor and visual average shift times across the 100 bootstrapping iterations for each participant. Coloured dots represent individual participants’ correlation values, the white line inside the grey box represents the mean, and the edges of the grey box represent the first (left) and third (right) quartiles.

The model was estimated using a maximum likelihood criterion, and the outputs of the model were reported as unstandardised regression coefficients with t-statistics. We used two-tailed tests and a 5% criterion for significance. Finally, we calculated the correlation (Pearson’s r) between the average visual and motor shift times across all bootstrapping iterations per participant. The resulting r values were statistically compared against a null hypothesis of 0 with a one-sample t-test (**Figure 4b**).

### EEG: relation to behaviour

We investigated the relation between the two behavioural dependent variables of interest (report error and RT) and the modulation of alpha- and mu/beta-frequency activity during the post-cue delay in informative trials separately for trials with a long and a short delay. For each participant and trial type (short and long), the median RT was estimated, and the trials were divided into those that were faster (fast) and those that were slower (slow) than the median RT. In parallel, the median report error was calculated. Trials were then separated into those with more (precise) or less accurate (imprecise) reports than the median error. Subsequently, we calculated and compared the contralateral-versus-ipsilateral alpha and mu/beta contrasts in fast vs slow trials and in precise vs imprecise trials (**Figure 5**).

**Figure 5.**
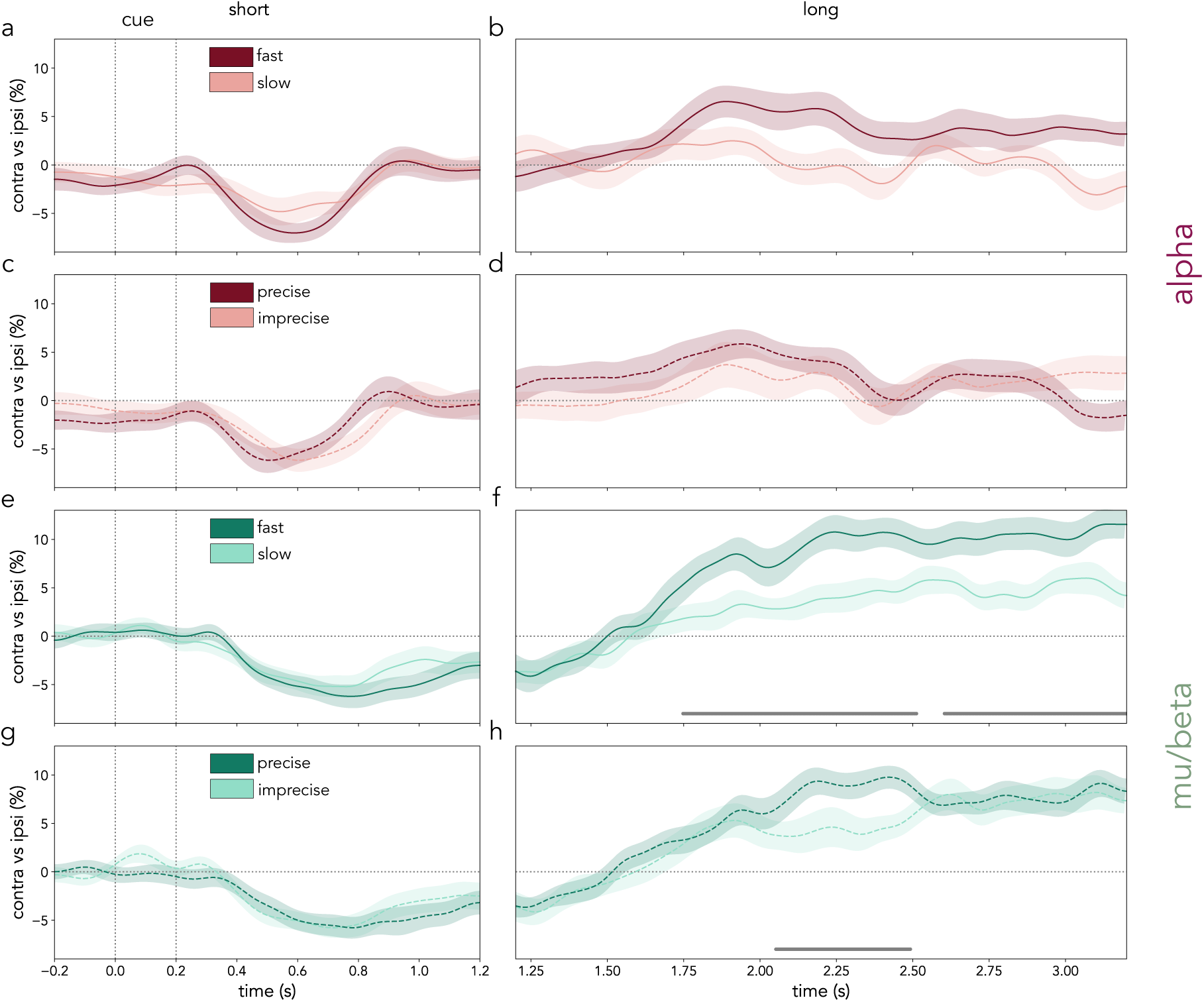
Relation between alpha and mu/beta modulations and behaviour. Alpha and mu/beta activity contralateral vs ipsilateral to the cued location/hand in fast vs slow and precise vs imprecise trials. a-d) Average alpha (8-12 Hz; burgundy) activity difference between contralateral and ipsilateral sensors to the cued location in trials categorised as fast (dark) or slow (light) based on a median split of RT (a,b) and in trials categorised as precise (dark) or imprecise (light) based on a median split of report error (c,d). The left column depicts short trials (a,c) and the right column shows long trials (b,d). e-h) Average mu/beta (8-30 Hz; green) activity difference between contralateral and ipsilateral sensors to the cued action in trials categorised as fast (dark) or slow (light) based on a median split of RT (e,f) and in trials categorised as precise (dark) or imprecise (light) based on a median split of report error (g,h). The left column depicts short trials (e,g) and the right column shows long trials (f,h). Cluster-based permutation significant time points of the contrast between the displayed time courses are indicated with horizontal grey lines. Shaded areas represent the SEM.

### Cluster-based permutation testing

Cluster-based permutation (Maris & Oostenveld, 2007) tested for statistical differences in the contralateral-versus-ipsilateral time-frequency spectra in informative vs noninformative trials and the alpha and mu/beta modulation time-courses. This statistical approach assumes the effects of interest are clustered across the relevant dimensions (e.g., time, time-frequency, space-time-frequency), making it suitable for testing the statistical significance of EEG activity patterns.

First, a mass-univariate t-test (two-sided, alpha = .05) was performed on the group-level contrasts, and the sum of all t-values in a cluster was defined as the cluster statistic for each given cluster. Next, the contrasts across the conditions of interest were randomly permuted (sign-flipped) 10,000 times for each participant. The largest clusters found under this null hypothesis were compared with the cluster statistics in the observed data. For each cluster, the proportion of permutations whose largest cluster exceeded the cluster identified in the observed data was calculated and the resulting p-value was estimated. Importantly, this approach circumvents the common problem of multiple comparisons by comparing distributions of the summary cluster statistics.

## Results

The present study investigated whether and how sensory and motor working-memory contents could be prioritised as a function of dynamically evolving expectations. With this aim, participants completed a visual-motor working-memory task in which retro-cues and dynamically changing expectations guided the prioritisation of sensory- and action-related working-memory contents. Crucially, this task enabled the independent tracking of sensory and action-related working-memory content prioritisation.

### Behavioural results

If participants prioritised each of the relevant working-memory contents at the expected times, their responses were hypothesised to be faster in both short and long informative trials compared to noninformative trials. A repeated-measures 2 x 2 ANOVA of RT with informativeness and duration as factors revealed a main effect of informativeness on RT (F(*1*,*29*) = 183.37, ***p < .001, **η**^2^ = .28), a main effect of duration on RT (F(*1*,*29*) = 101.17, ***p < .001, **η**^2^ = .09), and no interaction between the factors (F(*1*,*29*) = .78, p = .38, **η**^2^ < .000; **Figures 1b**; see also **Supplementary Tables 1 and 3**). Overall, participants were faster at responding to targets in informative trials than in noninformative trials, suggesting that both retro-cues and internally driven temporal expectations facilitated performance in this task. Participants were also faster in long trials compared to short trials, as expected from related foreperiod effects (Los, 2010; Niemi & Naatanen, 1981).

Report errors in this task were consistently low and insensitive to the informativeness of the retro-cue (F(*1*,*29*) = .003, p = .95, **η**^2^ < .001). A main effect of duration on error indicated smaller errors (higher accuracy) on short trials than long trials (F(*1*,*29*) = 24.25, ***p < .001, **η**^2^ = .02). The factors did not interact (F(*1*,*29*) = .02, p = .90, **η**^2^ < .001; **Figure 1c**; see also **Supplementary Tables 2 and 4**).

Although informativeness did not have a significant effect on report error, we confirmed that participants used orientation-related visual information to guide their responses by investigating the profile of the orientation reports. Namely, we found that participants’ responses varied systematically with the orientation of the probed bar (**Figure 1d**), rather than simply reflecting categorical left/right button presses. This confirms that participants were using the details of the visual representation in working memory to guide their reports.

### Alpha- and mu/beta-frequency activity modulation

Next, we turned to the pre-defined alpha- and mu/beta-frequency EEG activity during the post-cue delay. Based on previous studies (e.g., van Ede et al., 2019a), alpha activity contralateral to the selected item location was hypothesised to be modulated by the internal prioritisation of locations. Additionally, mu/beta activity contralateral to the prospective action hand was predicted to change together with prospective action selection. We utilised noninformative trials – in which participants could not reliably select sensory or response-related information before the probe – as a baseline for the selection signals in informative trials. Specifically, participant-wise cluster-based permutation analyses of the time-frequency spectra in the contralateral-versus-ipsilateral contrasts of visual and motor selection in informative vs noninformative trials confirmed our hypotheses (**Figure 2a,b**; see also **Supplementary Figure 2**).

A pronounced modulation in alpha activity contralateral vs ipsilateral to the selected item location was observed following the retro-cue in informative trials during the early part of the post-cue period, which contained both short and long trials (**Figure 2a**; first cluster: ***p < .001, time range: 0.27-0.8 s, frequency range: 5-25 Hz, max lateralisation time: 0.54 s, max lateralisation frequency: 11 Hz). Strikingly, in long trials, the side of relative alpha attenuation shifted after the short interval had elapsed (**Figure 2a**; second cluster: *p = .03, time range: 1.75-2.1 s, frequency range: 7-26 Hz, max lateralisation time: 1.93 s, max lateralisation frequency: 7 Hz). This “shift” in relative alpha attenuation from one hemisphere to the other, pointed to a sequential prioritisation of each working-memory item location in turn. Initially, the cued item that was expected to be probed early was prioritised (**Figure 2c**; first cluster: ***p < .001, time range: 0.31-0.81 s). With the passage of time, priority shifted to the location of the other item, which was expected to be probed late (**Figure 2c**; first cluster: ***p < .001; second cluster: *p = .04, time range: 1.7-2.04 s).

In addition to the modulation of contralateral alpha activity over occipital EEG channels, the prioritisation of stimulus locations in visual working memory has also been shown to be related to small biases in gaze position in the direction of internally selected items (e.g., Liu et al., 2022; van Ede et al., 2019b, 2021). The present study revealed a significant bias of gaze position towards the location of the cued item first, followed by a bias of gaze position towards the opposite location in long trials (**Supplementary Figure 4**).

In parallel to sensory prioritisation, a noticeable early reduction in mu/beta activity occurred in contralateral motor sensors according to the expected response hand in short trials (**Figure 2b**; first cluster: **p = .002, time range: 0.41-1.54 s, frequency range: 9-39 Hz, max lateralisation time: 1.24 s, max lateralisation frequency: 12 Hz). Strikingly, the side of mu/beta attenuation also shifted from one hemisphere to the opposite, mirroring the flexible prioritisation of successive action plans corresponding to the first and second items, respectively (**Figure 2b**; only long trials; second cluster: ***p < .001, time range: 1.76-3.2 s, frequency range: 7-39 Hz, max lateralisation time: 2.6 s, max lateralisation frequency: 36 Hz). These findings were confirmed on the average time course of mu/beta activity (**Figure 2d**; first cluster: **p = .005, time range: 0.38-1.42 s; second cluster: ***p < .001, time range: 1.71-3.2 s). Importantly, the present findings were not dependent upon EEG sensor choice as a larger set of visual and motor channels revealed an equivalent pattern of results (**Supplementary Figure 1**).

Similar to previous studies (e.g., van Ede et al., 2019a), the modulation of alpha activity linked to the prioritisation of item location had a lateralised occipital (visual) EEG topography (**Figure 2e**), while changes in mu/beta activity were predominantly confined to central (motor) EEG channels (**Figure 2f**). The prioritisation of the first and second location modulated alpha-frequency activity with comparable topographies. Similarly, the prioritisation of the first and second prospective actions had similar topographies and modulated a similar mu/beta frequency band.

### Temporal coupling between alpha and mu/beta modulation

Similar to what has been previously reported (van Ede et al., 2019a), visual inspection of alpha and mu/beta modulation showed they began largely together after the retro-cue (**Figure 3a**). Strikingly, the time courses of stimulus location and prospective action hand prioritisation subsequently diverged into distinct trajectories. Specifically, the prioritisation of location information was relatively short-lived in both early and late selection phases. Conversely, the prioritisation of the first and then second prospective hand actions was longer-lasting. These differences suggest that, even when sensory and action-related contents co-exist in working memory, their prioritisation can develop independently.

Thus, we investigated whether alpha and mu/beta activity modulations evolved in lockstep across the post-cue delay. First, we directly compared the time courses of sensory and motor prioritisation in long trials, expressed as a proportion of their respective maximum and minimum. We found statistically significant differences between the normalised sensory and response prioritisation time courses across participants (first cluster: ***p < .001; second cluster: ***p < .001; **Figure 3a**), which provided preliminary evidence that sensory and motor prioritisation did not co-evolve in lockstep. This finding was supported by an LMM of alpha and mu/beta modulation as a function of time and frequency band (alpha vs mu/beta) which revealed significant interactions between frequency band and time (**Supplementary Figure 3; Supplementary Table 5**).

Next, we investigated potential differences in the latency of the shift from prioritising one location and action to the other, as indexed by the alpha and mu/beta modulation time courses, respectively. With this aim, the average time at which the alpha (visual) and mu/beta (motor) time series crossed the y-axis (i.e. reversed their lateralisation) per participant was compared (**Figure 3b**; see *Methods* for details). A paired-samples t-test between the average *shift times* across participants revealed that the *visual shift*, quantified on the alpha time series, occurred significantly earlier than the *motor shift*, quantified on the mu/beta time series (t(*29*) = 4.3, ***p < .001, d = .87; **Figure 3c**). Interestingly, the average of the estimated *visual shift time* coincided approximately with the time of probe appearance in early trials (1.2 s from cue onset; M: 1.16 s; SD: .4 s). Alternatively, the *motor shift time* occurred at an average of 1.52 s (SD: .13 s) from cue onset, around 0.37 s (SD: 0.46 s) later than the visual shift time (**Figure 3c**).

As further evidence that sensory and motor prioritisation could evolve independently, we hypothesised that sensory and motor shift times would be uncorrelated. Therefore, we investigated the cross-trial co-variability in visual and motor shift timing using a bootstrapping procedure (see *Methods*). Across 100 iterations per participant, the motor shift time was not found to be a significant predictor of visual shift time in a LMM that included participant-related random effects (**β** = .02, t = .99, p = .32; **Figure 4a**). Next, we calculated the correlation (Pearson’s r) between visual and motor shift times across bootstrapping iterations per participant. A one-sample t-test revealed that the participant-wise correlation values were not significantly different from zero (t(*29*) = .24, p = .8, d = .1; **Figure 4b**). The absence of a correlation between visual and motor shift times further points to a functional decoupling of the alpha and mu/beta modulation time courses.

### Relation of alpha and mu/beta modulations to behaviour

Next, the relation between the behavioural variables of interest (RT and report error) and the alpha- and mu/beta-frequency activity patterns was investigated. To do so, trials were sorted based on a median split of performance (RT and error) per participant separately for short and long trials. Mu/beta activity modulations were found to relate to performance, as measured with RT and report error. Long informative trials with faster responses also displayed a stronger modulation of mu/beta-frequency activity in the hemisphere contralateral to the hand of the second prospective action (first cluster: **p = .002; second cluster: **p = .003; **Figure 5f**). Interestingly, despite not showing a general effect of informativeness on report error (**Figure 1c**), smaller errors (higher precision) in informative long trials were also related to a stronger mu/beta modulation (cluster: *p = .003; **Figure 5h**). No significant clusters in lateralised alpha activity were found in fast vs slow trials or precise vs imprecise trials (**Figure 5a-d**). Together, these results suggested that only modulations in mu/beta activity were tightly related to performance, in particular the modulations that signalled the change of action plan associated with the shift as time elapsed.

## Discussion

The present study reveals that both sensory and action-related contents that co-exist in working memory are flexibly and dynamically prioritised in a temporally structured but independent fashion.

In this task, both retro-cues and internal states related to changing temporal expectations successfully elicited attentional prioritisation of stimulus locations and associated actions in working memory. Across two sessions, response times were faster when responding to targets at their expected times. Importantly, the speed of responses to previously un-prioritised items was comparable to first-prioritised item responses, highlighting the flexibility and reversibility of internal attention (see De Vries et al., 2017, 2018; LaRocque et al., 2013; Lepsien & Nobre, 2007; Lewis-Peacock et al., 2012; Murray et al., 2013; Myers et al., 2018b; Rerko & Oberauer, 2013; Van Moorselaar et al., 2015). Perhaps due to participant over-training, accuracy was exceptionally high in this task and insensitive to attentional selection. Mirroring the behavioural results, the alpha and mu/beta modulation time courses uncovered that the prioritisation of both sensory- and action-related contents in working memory was flexible (i.e., it “shifted” from prioritising the first to the second location/action) and temporally tuned (i.e., both instances of selection of the relevant item/action were temporally specific).

The pattern of sensory (alpha) modulation in the present study replicates previous findings that the prioritisation of sensory contents in working memory is flexible (De Vries et al., 2017, 2018; LaRocque et al., 2013; Lewis-Peacock et al., 2012; Murray et al., 2013; Myers et al., 2018; Rerko & Oberauer, 2013; Van Moorselaar et al., 2015) and temporally tuned (van Ede et al., 2017; Zokaei et al., 2019). Consistent with previous studies, the short-lived modulation of alpha activity in the first and second selection events points to a transient prioritisation of item locations (see also De Vries et al., 2017, 2018; Gresch, et al., 2024a; Mok et al., 2016; Schneider et al., 2015, 2016). This has been suggested to reflect that spatial orienting helps activate working-memory contents into a prioritised state (Myers et al., 2017). In line with what has been previously reported (e.g., Foster et al., 2017; Kuo et al., 2009; Mok et al., 2016; Sligte et al., 2008; Sreenivasan et al., 2014), spatial selection in this task occurred even when the spatial arrangement of working-memory items was task-irrelevant. In contrast to other studies (e.g., van Ede et al., 2017), here, alpha-frequency activity during the memory delay was not systematically correlated with behaviour. We speculate that this could be related to the absence of a behavioural effect of selection on report error and the smaller magnitude of alpha modulation compared to mu/beta in this study.

To the best of our knowledge, the present study is the first to reveal the flexible, reversible, dynamic, and temporally tuned nature of proactive action-related prioritisation in working memory. Specifically, action-related contents that were deprioritised in the first instance could be re-prioritised later as indexed by a strong mu/beta modulation and behavioural benefits in long trials. Mu/beta modulation persisted well into the second part of the delay, after the time of early probe appearance had passed. This accords with the requirement to respond using the selected effector (left or right hand) at the end of the first or second selection period.

Importantly, the extent of mu/beta modulation in the second memory delay correlated with enhanced task performance. Mu/beta modulation was enhanced in faster trials as opposed to slower ones (see also Boettcher et al., 2021; Nasrawi et al., 2023), suggesting that the RT effects in this task may be reflective of differences in motor preparation. Interestingly, despite the insensitivity of report error to cue informativeness, mu/beta-frequency activity during the second part of the memory delay was stronger in precise than imprecise trials, highlighting the importance of motor states in the internal attention process. Together, these findings point to a key role for mu/beta activity in flexibly prioritising motor contents in working memory and highlight its value as a functionally relevant marker of internal attention.

A central finding of the current study with important theoretical consequences was the temporally dissociable modulation of sensory and action-related contents in working memory. Their flexible and dynamic prioritisation progressed in tandem but was uncoupled. In the first moments after the retro-cue, sensory and action-related prioritisation started together, consistent with previous findings (van Ede, et al. 2019a; see also Ester & Weese, 2023). However, in contrast to the transient nature of alpha modulation in this task, mu/beta activity showed a longer-lasting motor modulation. The present findings are consistent with previous studies suggesting that working-memory contents may undergo a change from a sensory to a motor code as the requirement to respond takes precedence in the task (González-García et al., 2020; Henderson et al., 2022; Kikumoto et al., 2022; Myers et al., 2017; Wallis et al., 2015).

Crucially, as expectations about the to-be-probed memory representation reversed, the sensory and action-related modulation signals evolved independently. The modulation of sensory and motor systems by internal attention, as measured with alpha and mu/beta activity modulations, did not co-evolve in lockstep. Even though they both showed a clear reversal with time, this reversal developed distinctly for both markers and its exact timing was uncorrelated. Together, these findings suggest that sensory and action-related dimensions of working-memory representations need not be intrinsically bound but instead can be dissociable and susceptible to separate modulatory processes. In turn, this highlights the multiplicity of modulatory processes that can operate on working-memory representations contemporaneously but independently (Nobre & van Ede, 2023; van Ede & Nobre, 2023).

Here, the modulation of location-related activity was transient, while the modulation of motor activity was longer-lasting. The patterns match the changes in when and for how long location- and action-related information was relevant to this task. More generally, we speculate that different attributes of working memory objects may be differentially modulated depending on changing expectations about their relevance. Relatedly, external attention has also been shown to differentially modulate spatial-sensory and motor systems when the purpose of the task emphasises perceptual demands vs speeded responses, respectively (van Ede et al., 2020).

In summary, the present study shows that sensory and action-related contents that co-exist in working memory are flexibly and dynamically prioritised and tuned to the moments when they are anticipated to become relevant. This points to a fine temporal organisation of internal attention according to external temporal regularities. Moreover, the prioritisation of sensory and action-related contents that co-exist in working memory was not continuously temporally coupled. This speaks to the relative independence of object dimensions in working memory and the multiplicity of flexible and dynamic modulatory processes that co-occur to prepare internal representations for adaptive behaviour.

## Acknowledgements

The authors would like to thank Lola Milton-Jenkins for her invaluable help with data collection, the members of the Brain and Cognition Lab for fruitful discussions, and all the participants who contributed their time and effort to participating in this study. This research was funded by a Wellcome Trust PhD Studentship (102170/Z/13/Z) I.E.A., a James S. McDonnell Foundation Understanding Human Cognition Collaborative Award (220020448) and a Wellcome Trust Senior Investigator Award (104571/Z/14/Z) to A.C.N, and an ERC Starting Grant from the European Research Council (MEMTICIPATION, 850636) to F.v.E. The Wellcome Centre for Integrative Neuroimaging is supported by core funding from the Wellcome Trust (203139/Z/16/Z). For the purpose of open access, the author has applied a CC BY public copyright licence to any Author Accepted Manuscript version arising from this submission.

## Supplementary Material

### Supplementary Methods

#### EEG: linear mixed effect model of alpha and mu/beta modulation

As a complementary analysis of the temporal coupling between the alpha and mu/beta modulation time courses, we fit a linear mixed-effects model (LMM; using the *lmer* function in the *lme4* R library) to the participant-averaged alpha and mu/beta modulation time-course as a function of time and frequency band according to the following formula:

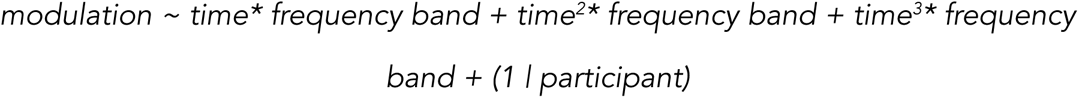

Here, modulation referred to the participant-averaged contra-vs-ipsi alpha and mu/beta activity modulation in units ranging from −1 (minimum) to 1 (maximum). The frequency-band variable had two levels referring to posterior alpha- (visual) or central mu/beta- (motor) activity respectively. Time represented the time from cue onset until probe onset in long trials centred around the hypothetical onset of the early probe (1.2 s from cue onset). We let the modulation vary as a function of linear, quadratic, and cubic time to capture the non-linear nature of the alpha and mu/beta modulation time courses. Moreover, we let linear, quadratic, and cubic time interact with frequency-band (alpha or mu/beta) to capture linear and non-linear differences in the temporal unfolding of visual and motor prioritisation. Finally, we included participant-related random effects in the model. We used the alpha frequency band as the reference contrast. All the coefficients in **Supplementary Table 5** ought to be interpreted accordingly. The model was estimated using a maximum likelihood criterion, and the outputs of the model were reported as unstandardised regression coefficients with t-statistics and 99% confidence intervals (**Supplementary Figure 3; Supplementary Table 5**). We used two-tailed tests and a 5% criterion for significance.

#### Eye-tracking data analyses

In session 2 (EEG), bilateral eye position was continuously monitored with an eye-tracking device at a sampling rate of 1000 Hz (Eyelink 1000, SR-Research Ltd., Ottawa, Ontario, Canada). Participants performed an eye-tracking calibration task before blocks 1, 6, and 11 of the experiment. Additional calibration tasks were performed if ocular drift was noticed between these moments. Calibration did not work for three participants and eye-tracking data could not be collected. One participant was excluded who did not meet the behavioural inclusion criteria. Additionally, three other participants were excluded because the eye-tracking signal was lost for more than 50% of the trials. In total, eye-tracking data from 23 participants were analysed.

The eye-tracking signal was pre-processed following the steps detailed in related studies (e.g., Draschkow et al., 2022; Gresch et al., 2024b; van Ede et al., 2019b). First, the acquired edf files were converted into the asc format using EDFConverter (SR-Research Ltd.). All subsequent analyses were performed using R studio (RStudio Team, 2020), the *eyelinker* R library, and custom-made scripts.

Eye blinks were identified and the signal +/-100 ms around each blink was discarded following the guidelines from the Eyelink manual (Eyelink 1000, SR-Research Ltd., Ottawa, Ontario, Canada). Subsequently, data from the left and right eye were averaged yielding one time course for eye movements along the horizontal axis (x-position) and another along the vertical axis (y-position). Only the horizontal gaze position was analysed further, as the bars were positioned on the screen along the horizontal axis.

Gaze position data were downsampled to 250 Hz. Next, data were epoched from 500 ms before to 3400 ms after cue onset in long trials and between −500 and 1400 ms in short trials. Trials with eye movements exceeding half the distance to the bars (96 pixels) were removed from further analyses (see also Gresch et al., 2024b). Additionally, trials with RTs below 100 ms or with incorrect responses were discarded from further analyses. An average of 13.05% of trials (SD: 11.15%) were excluded. The time course of the horizontal gaze position in each trial was smoothed using a Gaussian kernel with a standard deviation of 40 ms. Subsequently, the epochs were cropped between 200 ms before and 1200 ms (short) or 3200 ms (long) after the cue to remove any smoothing-related edge artefacts. Finally, the average baseline activity (−200 to 0 ms), was subtracted from each epoch.

Leftward and rightward gaze-position time courses were compared between trials where the left or the right item was cued (**Supplementary Figure 4a**). “Towardness” was calculated as the average between the gaze position in trials where the right item was prioritised and the sign-flipped gaze position in left-item trials. Thus, towardness indicated the extent of biases in gaze position towards the prioritised item location. Cluster-based permutation tested for statistically significant differences between the different time courses.

### Supplementary Figures

**Supplementary Figure 1.**
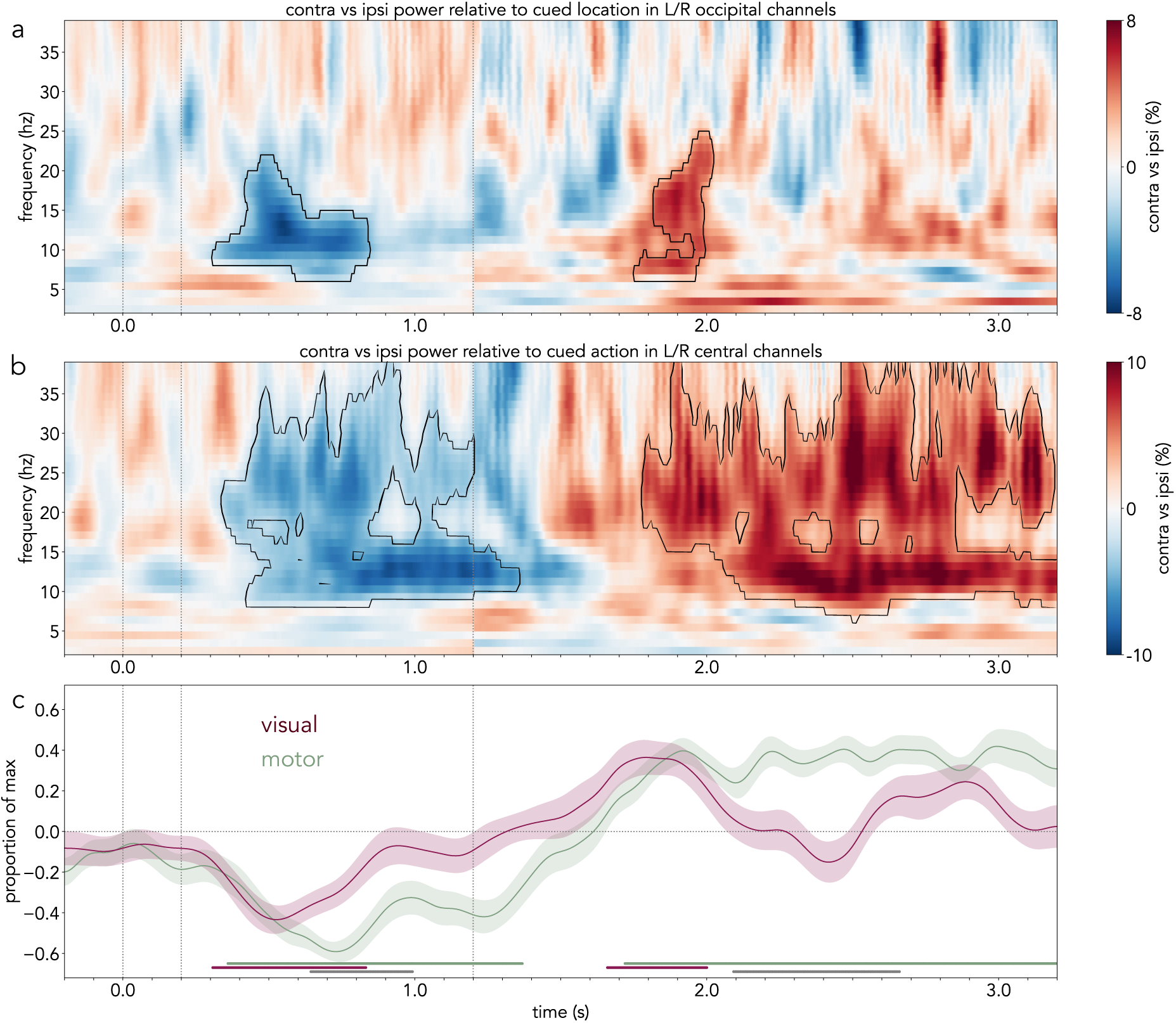
Lateralised, frequency-specific EEG activity locked to cue onset in informative trials in a cluster of visual/motor EEG electrodes. a) Contrast between EEG time-frequency activity contralateral vs ipsilateral to the cued bar location in two clusters of lateralised occipital sensors (L: O1, PO7, PO3; R: O2, PO8, PO4) divided by summed contralateral and ipsilateral activity (expressed as a percentage) in informative vs noninformative trials. Black outline indicates significant clusters. b) Contrast between EEG time-frequency activity contralateral vs ipsilateral to the cued prospective action in two clusters of lateralised central sensors (L: C1, C3, CP1, CP3; R: C2, C4, CP2, CP4) divided by summed contralateral and ipsilateral activity (expressed as a percentage) in informative vs noninformative trials. Black outline indicates statistically significant clusters. c) Average alpha (8-12 hz) activity difference between contralateral and ipsilateral sensors to the cued location across participants (burgundy) in informative trials. Average mu/beta (8-30 hz) activity between contralateral and ipsilateral sensors to the cued action across participants (green) in informative trials. Shaded areas represent the SEM and cluster-based permutation-corrected significant time points are indicated with horizontal lines (burgundy: alpha vs null; green: mu/beta vs null; grey: alpha vs mu/beta). The first part of the time-frequency spectra in panels a and b and of the time course in c (−0.2-1.2 s) corresponds to the average of short and long trials, and the second part (1.2-3.2 s) corresponds to long trials only. The vertical dotted lines represent (from left to right) the onset (0 s) and offset (0.2 s) of the retro-cue and the time of probe appearance in early trials (1.2 s). For comparison purposes, the time-frequency spectra are plotted on the same scale as Figure 2.

**Supplementary Figure 2.**
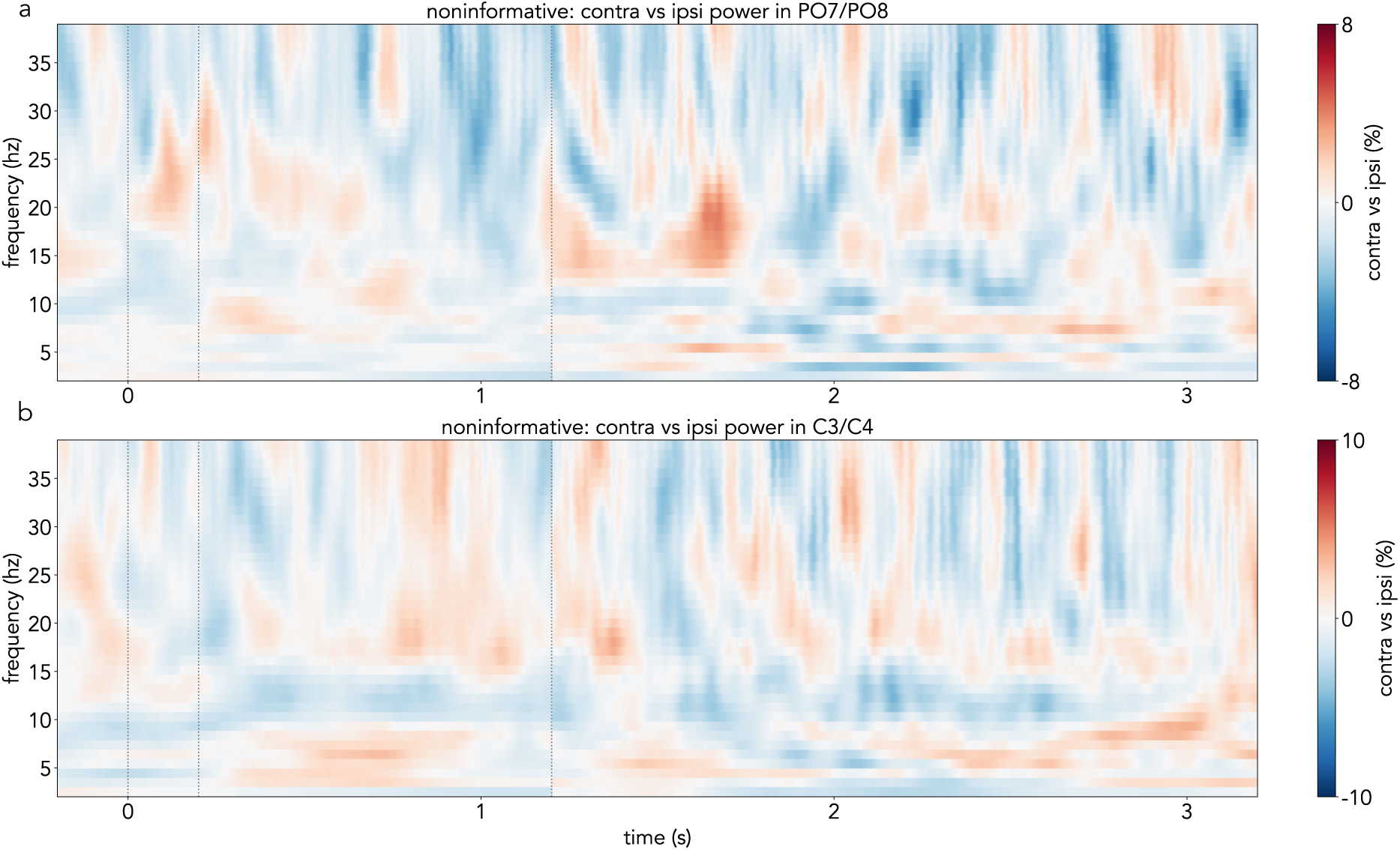
Lateralised, frequency-specific EEG activity locked to cue onset in noninformative trials, where no location/action is systematically prioritised. a) Contrast between EEG time-frequency activity contralateral vs ipsilateral to the cued bar location (none) in occipital sensors (PO7, PO8) divided by summed contralateral and ipsilateral activity and expressed as a percentage in noninformative trials. b) Contrast between EEG time-frequency activity contralateral vs ipsilateral to the cued prospective action (none) in central sensors (C3, C4) divided by summed contralateral and ipsilateral activity and expressed as a percentage in noninformative trials. The first part of the time-frequency spectra (−0.2-1.2 s) corresponds to the average of short and long trials, and the second part (1.2-3.2 s) corresponds to long trials only. The vertical dotted lines represent (from left to right) the onset (0 s) and offset (0.2 s) of the noninformative cue and the time of probe appearance in early trials (1.2 s). No significant clusters were found. For comparison purposes, the time-frequency spectra are plotted on the same scale as Figure 2.

**Supplementary Figure 3.**
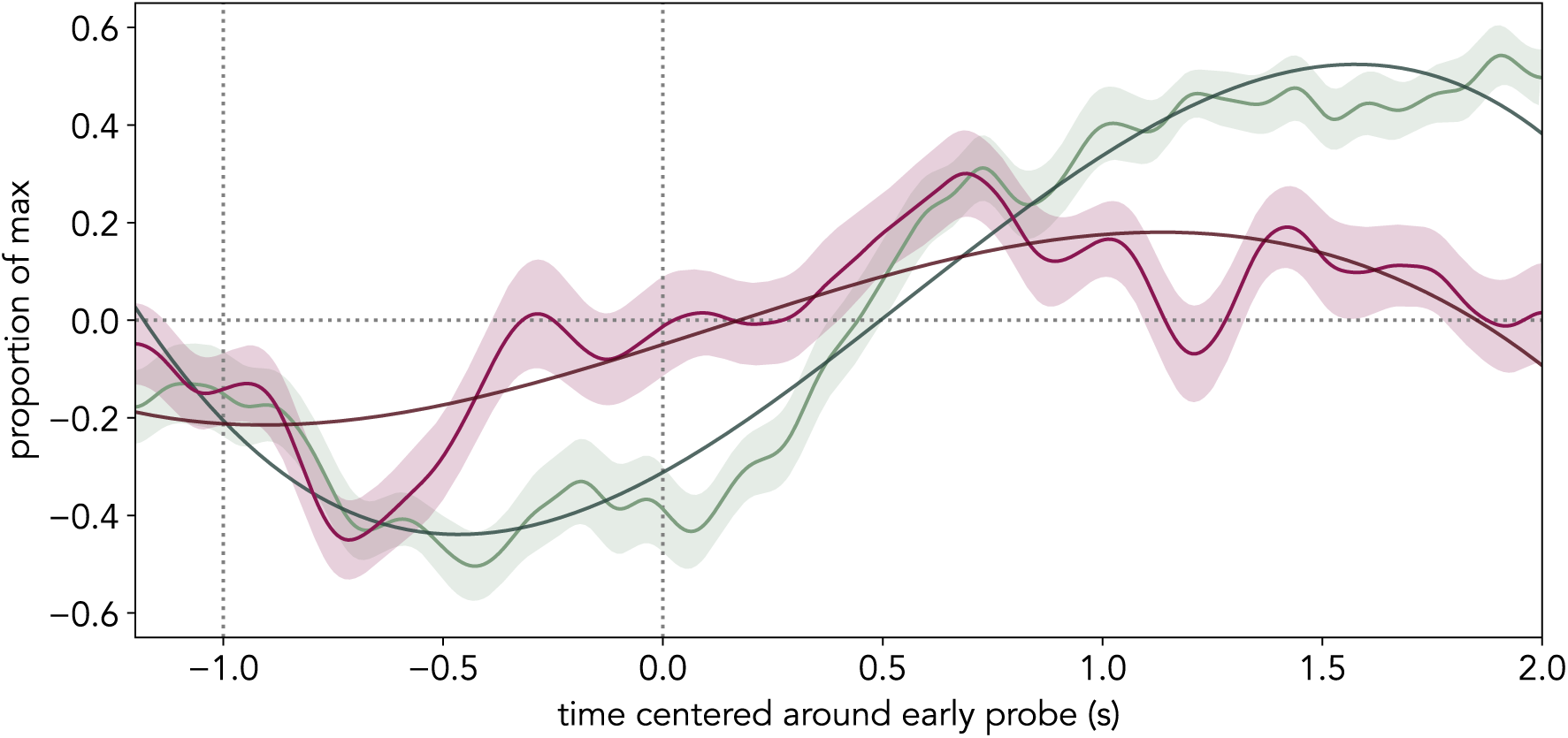
Linear mixed-effect model of the alpha and mu/beta modulation time courses as a function of time. Average alpha (8-12 Hz) activity difference between contralateral and ipsilateral sensors to the cued location in informative long trials (burgundy) and average mu/beta (8-30 Hz) activity between contralateral and ipsilateral sensors to the cued action (green) in informative long trials, both scaled to range between −1 and 1 and centred around the time of early probe onset (1.2 s). The dark lines depict the alpha and mu/beta modulation values as estimated with the linear mixed-effects model (see *Supplementary Methods*). Shaded areas represent the SEM. The vertical dotted lines represent (from left to right) the offset (−1 s) of the retro-cue and the time of probe appearance in early trials (0 s).

**Supplementary Figure 4.**
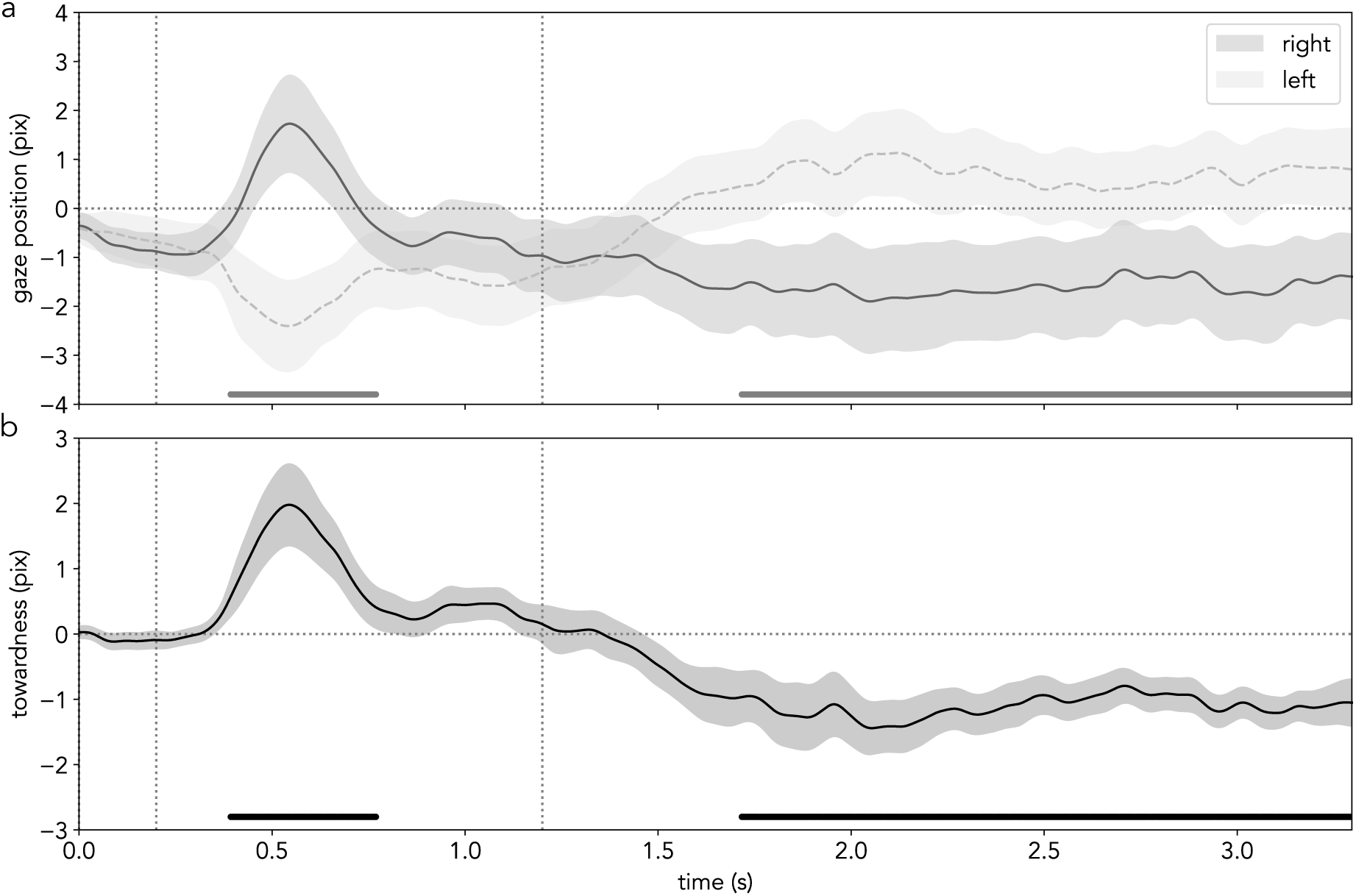
Horizontal gaze position as a function of cued item location locked to retro-cue onset in informative trials. a) Participant-averaged horizontal gaze position (in pixels) when an item on the left (dotted line) vs on the right (solid line) was cued locked to cue onset in informative trials (first cluster: *p = .01; second cluster: ***p < .001). b) Participant-averaged towardness (metric which collapses across left and right cued items) locked to cue onset in informative trials (black; first cluster: **p = .009; second cluster: *p = .02). Shaded areas represent the SEM, and vertical dotted lines represent (from left to right) cue offset and time of probe appearance in early trials. Cluster-permutation significant time points are indicated with horizontal lines at the bottom of the plots. The first part of the time courses (0-1.2 s) is the average across both short and long trials, and the second part (1.2-3.2) averages across long trials only.

### Supplementary Tables

**Supplementary Table 1.** The mean (M) and standard deviation (SD) of reaction time (RT) for different trial types in the behavioural and eeg sessions.

| *session* | *informativeness* | *duration* | *M* | *SD* |
| --- | --- | --- | --- | --- |
| behav | inf | long | 0.51 | 0.05 |
| behav | inf | short | 0.59 | 0.05 |
| behav | noninf | long | 0.66 | 0.04 |
| behav | noninf | short | 0.72 | 0.04 |
| eeg | inf | long | 0.51 | 0.05 |
| eeg | inf | short | 0.59 | 0.05 |
| eeg | noninf | long | 0.66 | 0.04 |
| eeg | noninf | short | 0.72 | 0.04 |

**Supplementary Table 2.** The mean (M) and standard deviation (SD) of report error for different trial types in the behavioural and eeg sessions.

| *session* | *informativeness* | *duration* | *M* | *SD* |
| --- | --- | --- | --- | --- |
| behav | inf | long | 9.34 | 1.15 |
| behav | inf | short | 9.26 | 0.86 |
| behav | noninf | long | 9.66 | 0.77 |
| behav | noninf | short | 9.15 | 0.88 |
| eeg | inf | long | 9.43 | 0.79 |
| eeg | inf | short | 8.72 | 0.68 |
| eeg | noninf | long | 9.24 | 0.70 |
| eeg | noninf | short | 8.79 | 0.54 |

**Supplementary Table 3.** ANOVA results for RT in the behavioural and eeg sessions. Ges reflects η^2^.

| *session* | *factor* | *df* | *F* | *p* | *ges* |
| --- | --- | --- | --- | --- | --- |
| behav | informativeness | 28 | 135.27 | <0.001 | 0.25 |
| behav | duration | 28 | 96.25 | <0.001 | 0.10 |
| behav | informativeness:duration | 28 | 1.91 | 0.18 | 0.00 |
| eeg | informativeness | 29 | 188.42 | <0.001 | 0.26 |
| eeg | duration | 29 | 78.71 | <0.001 | 0.07 |
| eeg | informativeness:duration | 29 | 0.11 | 0.74 | 0.00 |

**Supplementary Table 4.** ANOVA results for report error in the behavioural and eeg sessions. Ges reflects η^2^.

| *session* | *factor* | *df* | *F* | *p* | *ges* |
| --- | --- | --- | --- | --- | --- |
| behav | informativeness | 28 | 0.28 | 0.6 | 0.00 |
| behav | duration | 28 | 3.88 | 0.06 | 0.00 |
| behav | informativeness:duration | 28 | 1.61 | 0.21 | 0.00 |
| eeg | informativeness | 29 | 0.23 | 0.64 | 0.00 |
| eeg | duration | 29 | 20.72 | <0.001 | 0.03 |
| eeg | informativeness:duration | 29 | 1.49 | 0.23 | 0.00 |

**Supplementary Table 5.** The coefficients of the fixed and random effects of the LMM which modelled alpha and mu/beta modulation as a function of frequency-band and linear, quadratic, and cubic time (see *Supplementary Methods*).

| *Predictors* | *modulation* |  |  |
| --- | --- | --- | --- |
|  | *Estimates* | *CI* | *p* |
| (Intercept) | -0.05 | -0.09 - -0.01 | **0.006** |
| freqB | -0.26 | -0.28 - -0.25 | **<0.001** |
| time_1 | 0.29 | 0.27 - 0.30 | **<0.001** |
| time_2 | 0.03 | 0.02 - 0.04 | **<0.001** |
| time_3 | -0.09 | -0.10 - -0.08 | **<0.001** |
| freqB:time_1 | 0.21 | 0.19 - 0.23 | **<0.001** |
| freqB:time_2 | 0.35 | 0.33 - 0.36 | **<0.001** |
| freqB:time_3 | -0.13 | -0.15 - -0.12 | **<0.001** |
| **Random Effects** |  |  |  |
| $\sigma^2$ | | 0.19 | |
| T00_snum |  | 0.01 |  |
| ICC |  | 0.04 |  |
| N_snum |  | 30 |  |
| Observations |  | 48060 |  |
| Marginal R<sup>2</sup> / Conditional R<sup>2</sup> |  | 0.266 / 0.298 |  |

## Notes

### Competing Interest Statement

The authors have declared no competing interest.

### Summary of Updates

Figure 5 revised. Labels in panels a and c were reversed and labels in panels e and g were also reversed. Updated figure added to manuscript pdf.

